# Development of Microsatellite Markers for a Soricid Water Shrew *Chimarrogale platycephalus*, and the Success of Individual Identification Using It

**DOI:** 10.1101/2020.05.21.108324

**Authors:** Haruka Yamazaki, Tomohiro Sekiya, Shun Nagayama, Kei Hirasawa, Keita Tokura, Akio Sasaki, Hidetaka Ichiyanagi, Koji Tojo

## Abstract

The soricid water shrew, *Chimarrogale platycephalus*, is a mammal species endemic to the Japanese Islands. They inhabit the islands of Honshu and Kyushu, and are considered to be extinct in Shikoku. Information on this water shrew from Honshu and Kyushu is also scarce, and *C. platycephalus* is registered on the Japanese Government’s red list as an endangered species. Almost all study areas such as regarding their ethology, ecology, also their phylogenetics are lacking. The delay in these foundational studies is due to difficulties related to them being both nocturnal and aquatic. In order to study of *C. platycephalus*, it is essential to conduct field research in mountain streams at night under such circumstances. In response to such difficult circumstances, we established a genetic analysis method using the feces of *C. platycephalus*, and as a result the accumulation of data on their phylogenetic evolution and phylogeography has increased which has improved our understanding of this species. Furthermore in this study, development of microsatellite markers was carried out, and a total of 21 loci marker analyses were performed. In addition, in order to confirm the credibility of these 21 microsatellite markers, as being able to validly differentiate individuals, all markers were examined using the fecal and tissues specimens from verified separately reared individuals (12 individuals) with the known history of having been reared in an aquarium. As a result, it was revealed that if analyses of the 12 loci were possible, individual differentiation with 100% accuracy was possible. The development of microsatellite markers in this study, and the establishment of individual identification methods by means of this will be expected to greatly contribute to future ecological and ethological research, population genetics and biogeographical research of the water shrew, *C. platycephalus*.

## INTRODUCTION

A freshwater soricid shrew *Chimarrogale platycephalus* (Soricomorpha, Soricidae) endemic to Japan is recorded on only the two main islands, i.e., Honshu and Kyushu. There are no records of *C. platycephalus* in Hokkaido and Shikoku. The island of Hokkaido, which is located north of the Tsugaru Strait, often functions as a distribution boundary for many organisms, and is thus considered to be situated outside of this species’ distribution area. On the other hand for Shikoku, for which this species does not have any distribution records, it is suggested that this species may have become extinct (Abe 2003).

*Chimarrogale platycephalus* inhabits mountain streams, preying on benthos such as aquatic insects and freshwater crabs, and also freshwater mountain fish (i.e, salmonid *Salvelinus leucomaenis* and *Oncorhynchus masou*) (Fig. S1). This is an ecologically important position, being one of the top predators in mountain stream ecosystems. Even in Honshu and Kyushu, they are ranked within one of the categories on the local government’s red list, and are a target for conservation.

As it is extremely vulnerable to stress due to capture by trapping, investigation and/or research on its behavior/ecology is hindered. Under such circumstances, we tried to establish methods and techniques for their genetic structure analyses utilizing their feces, and we were able to successfully obtain concise results (Sekiya et al. 2017). So far, we have been conducting biogeographic studies targeting wide areas across Honshu and Shikoku. By evaluating these analysis results, it is suggested that the analysis method utilizing mitochondrial DNA Cyt b region is almost universally established. Currently, by means of increasing the number of regional groups being analyzed, we are developing research outcomes in the field of biological phylogeography, which is yielding higher accuracy results.

In addition, there are many points of interests with respect to this species, as follows. How large is their behavior and foraging on the individual level? Is there a difference in the behavior range between males and females? What kind of family structure do they maintain? Is there territorial behavior? To assist in such studies, we attempted to develop microsatellite markers that enable identification of individuals of *C. platycephalus*. As a result, good microsatellite markers were able to be obtained (21 genetic loci). In addition, it was confirmed that individual identification of sufficient precision is possible by combining these microsatellite markers.

In this study, we conducted experimental verification of twelve previously identified individuals, which were subsequently reared separately. We then in a blind test extracted and purified their total genomic DNA from their feces, and performed fragment analysis using the microsatellite markers we had previously developed. We confirmed whether it is possible to identify individuals being reared even under masked conditions by their feces. As a result, the correct identification rate achieved for the feces of the twelve individuals was 100%. These reared individuals included individuals collected from closely located sites within the same mountain stream. Although these individuals were genetically closely related to each other, it was still possible to maintain a correct answer rate of 100%. Thereby, we have demonstrated that this technique makes it possible to identify individuals at quite a high level.

In addition, in the analysis of fecal samples taken from the field, interesting cases were also detected in which multiple individuals share the same “dropping” site. In this article, we will detail the development of a robust process for selection of suitable microsatellite markers and introduce some practical examples for the use of these markers.

## MATERIALS AND METHODS

### 1. Microsatellite (SSR) marker analysis using the NGS (Next Generation Sequencer)

#### 1.1. Materials

A tissue specimen (not feces specimens) of the freshwater soricid shrew *Chimarrogale platycephalus* captured in Shizuoka Prefecture was used, in order to extract its high purity total genomic DNA. The specimen collected within a series of biodiversity surveys in Shizuoka Prefecture had one part of it (its tail) dissected. This specimen was stored frozen in 99% ethanol.

#### 1.2. Total genomic DNA extraction from the *Chimarrogale platycephalus* tissue

The DNeasy Blood & Tissue Kit (QIAGEN, Hilden) was used to extract the total DNA extraction from the *C. platycephalus* tissue. Using the tissue from the tail as a specimen (5.0 mg), extraction and purification was carried out largely according to the protocol recommended by the product manufacturer. This specimen (tissue sample) was placed in 180 μl ATL buffer, subsequently 20 μl of proteinase K was added, and then the DNA lysate was obtained after the cells disaggregated after stirring for about 24 hours at 55 °C. 100 μl of AL buffer and 200 μl of 100% ethanol were added to this DNA lysate, subsequently by passing the solution through a special filter the DNA only was collected on the filter. Thereafter, 500 μl of the AW 1 (washing solution) was added and a centrifuge was used to wash impurities through, followed by adding 500 μl of the AW 2 (washing solution) and centrifuging again to remove impurities. Finally, the DNA adsorbed on the filter was eluted with 200 μl of AE buffer (QIAGEN, Hilden) to obtain the DNA solution.

Using the spectrophotometer, Nano Vue Plus (GE Healthcare, Buckinghamshire), the concentration was adjusted using an eluate so that the concentration of the total genomic DNA was about 20 ng/μl, and thereafter the template was stored at 4 °C.

#### 1.3. NGS analysis, and the search for microsatellite sites

Using the extracted and purified total genomic DNA of *C. platycephalus*, analysis was performed using the NGS (Next Generation Sequencer) as described in detail below, and from the obtained sequence data, the extraction of microsatellite containing sequences was carried out. For preparation of a sequence library, the total genomic DNA of *C. platycephalus* was treated with the restriction enzyme EcoRI and ligated with P1 Adapters. After that, shear and end repair were performed at random, P2 Adapters were newly ligated to further fragments. By performing PCR using the both P1 and P2 Adapter sequences as primer set, only DNA fragments with P1 and P2 adapters added to both ends were PCR amplified. Among the fragments amplified by such PCR, those having a size of 300-500 bp were selected. By the quantitative real-time PCR (qPCR), libraries of the sequences of the genomic DNA of *C. platycephalus* with an appropriate insert size with effective concentrations of 2 nM or more were obtained by paired end sequences using the NGS, HiSeq X Ten (Illumina, California). Sequence assembly was carried out using the Velvet Optimizer (Zerbino and Birney 2008), after excluding low quality data by filtering. Primer design for detecting simple sequence repeats from the assembled contigs obtained of *C. platycephalus* genomic DNA and amplifying all detected repeat sequences was conducted using the software Primer 3 (Rozen and Skaletsky 2000).

#### 1.4. Selection of primers

Based on the obtained sequence data, suitable primers for our microsatellite analyses were selected according to the following four conditions.

##### 1.4.1. Number of repeat units

First, they were selected based on the number of microsatellite repetitions. The more repetitions, the more polymorphisms are likely to occur (Weber 1990, Ellegren 2000, Petit et al. 2005, Kelkar et al. 2008), but allele dropout increases in repeating sequences (Kirov et al. 2000, Buchan et al. 2005), which may cause problems such as an increase in stuttering (Hoffman and Amos 2005) that results in multiple peaks around the correct allele. According to Asch et al. (2010), it is recommended to select as target instances of 11-16 repeats within trinucleotides. In consideration of these things, we selected instances of over 8 repetitions in this study.

##### 1.4.2. Repeat unit size

Next, selection was based on the number of bases present in the repeating unit of microsatellites. In general, the number of different bases within microsatellites is 1 to 6 bases, and the shorter the allele size range, the easier it is for Multiplex PCR to analyze multiple loci at a time. However, on the other hand, stutter peaks tend to occur in short repeat units (Chambers and MacAvoy 2000), which increases the possibility of erroneous identification of alleles (Levinson and Gutman 1987, Meldgaard and Morling 1997). On the contrary, since it is difficult to perform the Multiplex PCR when long microsatellite repeat units of such as 5 or 6 bases are repeated, microsatellites of 2 to 4 bases were selected in this study.

##### 1.4.3. The Tm value of the primer

The specificity of PCR decreases when the Tm value is lowered, and the possibility of amplifying the DNA of an organism other than the shrew, *C. platycephalus* and a region other than the target region is increased. Therefore, in this study, the Tm value was set to around 60 °C. Furthermore, in Multiplex PCR which analyzes multiple loci at the same time, since multiple primer sets are used at the same time, the Tm values ware limited to within a range around 60 °C (i.e., between 60.0 and 62.6 °C), in order to regulate the amplification efficiency of all of the primer sets.

##### 1.4.4. Specificity of the DNA of *C. platycephalus*

These sequences selected as primer candidates were then searched using the software, Primer BLAST (Ye et al. 2012), and were thereby checked for homology with other organisms. In addition to fish and aquatic insects that *C. platycephalus* consume as foods, primers with high homology to human sequences were excluded to avoid the risk of contamination.

#### 1.5. Amplification of DNA fragments by PCR

The PCR was finally carried out using the primers and PCR conditions selected as described above. Successful DNA amplification was confirmed by electrophoresis the PCR product. The extracted and purified total genomic DNA from a tissue specimen of *C. platycephalus* was used as the template DNA for PCR. Regarding the PCR of this tissue specimen, KOD FX neo (TOYOBO, Osaka) was used as a DNA polymerase. At that point, four fluorescent markers for detecting the microsatellite sequences were added, i.e., the FAM, PET, NED and HEX sets (Applied Biosystems, Tokyo). As for PCR, the thermal cycler PC 350 (Astec, Fukuoka) was used. PCR conditions were as per the following. Firstly pre-heating for 2 minutes at 94 °C, followed by 28 cycles of “denaturation at 98 °C for 10 seconds, annealing at 60 °C for 30 seconds and extension at 68 °C for 30 seconds, per cycle”, with the final extension was carried out at 68 °C for 5 minutes. After this PCR, electrophoresis was carried out to confirm the presence or absence of copy amplification of the target DNA fragments.

#### 1.6. Sequencing

All of the primer sets in which DNA amplification was confirmed by electrophoresis were used, with which acual sequencing was carried out. Hi-Di Formamide (Applied Biosystems, California) and 500 liz size standard (Applied Biosystems, California) were added to the PCR product and sequencing was conducted using the automated DNA Sequencer (ABI 3130xl Genetic Analyzer; Perkin Elmer/Applied Biosystems). Using the software Genemapper ver. 4.1 (Applied Biosystems, California), the size of the DNA sequences was confirmed, alleles were determined, and genotyping was performed.

### 2. Establishment an individual identification method using microsatellite (SSR) markers

#### 2.1. Materials

The feces of twelve individuals of *C. platycephalus* separately reared in an aquarium (Aquamarine Inawashiro Kingfishers Aquarium, Inawashiro, Fukushima Prefecuture, Japan) were used as samples for gene analysis (Fig. 1). Feces were collected during the period from December 7, 2017 to December 20, 2017 and January 10, 2019 to Feburuary 18, 2019. Two to four samples of feces per individual were collected on different days, and a total of 29 samples were obtained (Fig. 2). Approximately half the volume of each feces sample (100-200 mg) was immersed in 3 ml of Inhibit EX Buffer (QIAGEN, Hilden) and stored at room temperature, and then mixed with Voltex mixer just before extracting its total genomic DNA. In addition, tissue of six individuals which died under rearing condition between 2017 and 2019 were collected and added to this analysis.

**Fig. 1.**
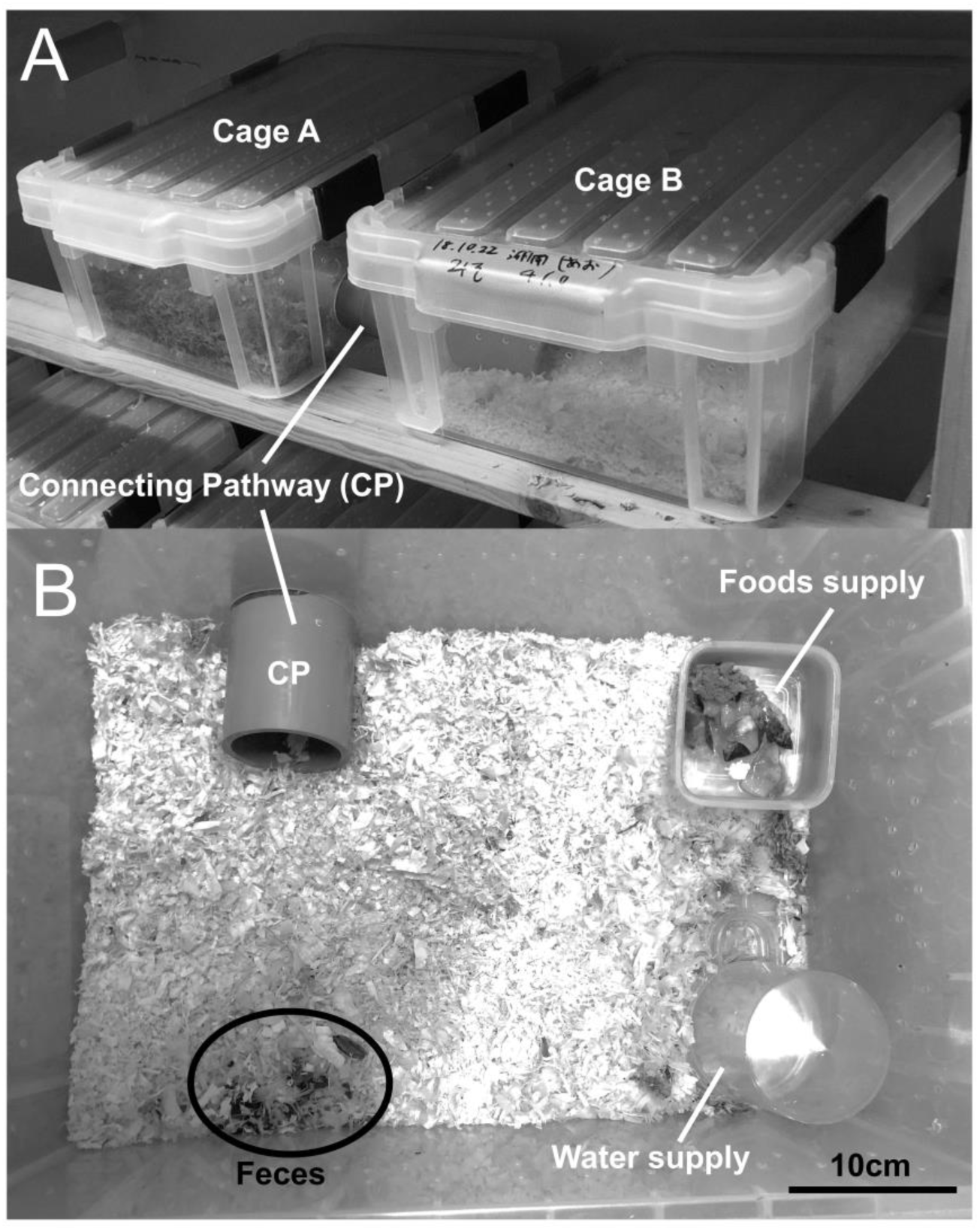
State of *Chimarrogale platycephalus* individuals reared, and excreted feces in their cage. The bottom of the rearing cage is covered with sawdust. Because they are extremely vulnerable to stress, the two rearing cases they are kept in are connected by a poly-vinyl chloride (PVC) tube so that they can move freely between each case (A). The lid was removed from one of the connected rearing cases, and photographed from the top (B). *1: PVC tube connecting two breeding cases. *2: Feeding container and food. In this breeding case, feces were excreted in the circle.

**Fig. 2.**
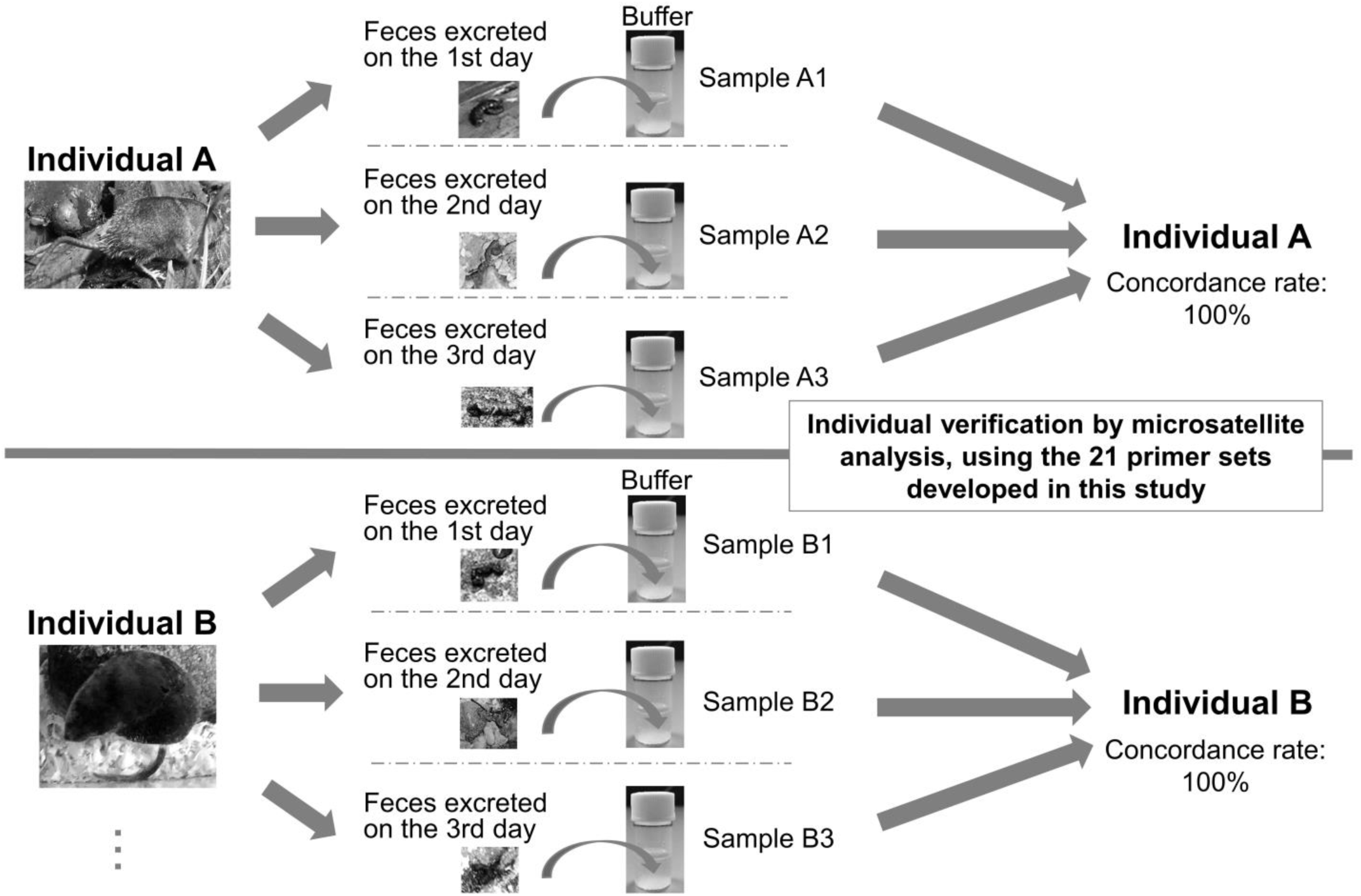
Flow of microsatellite analysis from *Chimarrogale platycephalus* fecal samples and the scheme of the experiments repeated. As for the experiments repeated, feces collected on different days were used.

#### 2.2. Total genomic DNA extraction and purification and genetic analyses from *C. platycephalus* feces

QIAamp Fast DNA Stool Mini Kit (QIAGEN, Hilden) was used for the total genomic DNA extraction and purification from fecal samples of *C. platycephalus*. Basically, we extracted and purified the total genomic DNA according to the methods developed in our previous research (Sekiya et al. 2017), which was a slight improvement on the protocol recommended by the product manufacturer.

Impurities were removed by centrifuging with Inhibit EX Buffer and the feces-turbid solution. Proteinase K (25 μl) and AL buffer (600 μl) were added to 600 μl of the resultant supernatant after centrifuging, and the mixture was left to stand at 70 °C for 10 minutes to allow the cells to disaggregate and obtain a lysate containing DNA. 600 μl of 100% ethanol was added to this lysate, and DNA was collected on the filter by passing the solution through a special filter. The washing solutions were added and centrifuged twice to remove impurities. Finally, the total genomic DNA was extracted and purified by eluting the DNA from the filter using 100 μl of ATE buffer. The concentration of total genomic DNA extracted in this way was measured using the spectrophotometer, Nano Vue Plus (GE Healthcare, Buckinghamshire), and then stored at 4 °C. In this case, with respect to the total genomic DNA measured to be at a concentration of 20 ng/μl or more, it was prepared using a dilution buffer (i.e., ATE buffer) so as to achive a concentration of about 20 ng/μl.

Regarding the experiments after the total genomic DNA extraction/purification, identical gene analyses were performed on the feces extracted ANS samples in the same manner as those performed after DNA extraction and purification from the tissue sample. However, since fecal samples contain substances that inhibit DNA analysis, PCR was carried out using the MightyAmp™ DNA Polymerase ver. 3 (Takara, Shiga), which is capable of resistance/nonspecific amplification and smear suppression against such PCR-inhibiting substances. In addition, bovine serum albumin (BSA Bovine serum albumin), which is a PCR stabilizer, was added. PCR conditions were as per the following. Firstly pre-heating for 2 minutes at 98 °C, followed by 35 to 38 cycles of “denaturation at 98 °C for 10 seconds, annealing at 60 °C for 30 seconds and extension at 68 °C for 1 minutes, per cycle”, with the final extension was carried out at 68 °C for 7 minutes. Since these analyses are based upon a trace amount of DNA collected from feces, at least two repetitions of each experiment were carried out, in order to ensure the reliability of the results. When two samples were not able to confirm PCR copy amplification of DNA fragments the results were excluded from the analyses. When the first and second results differed or the peak was small, verification was carried out by further repeating the analysis. If the analysis result was not stable after multiple iterative experiments, it was excluded from the subsequent analyses. Using the software Genemapper ver. 4.1 (Applied Biosystems, California), the size of the DNA sequences were confirmed, alleles were determined, and genotyping was performed.

#### 2.3. Analysis by GENECAP

In order to statistically verify whether multiple fecal samples from the same individual were confirmed to be feces of the same individual by this microsatellite (SSR) analysis, the software GENECAP version 1.4 (Wilberg and Dreher. 2004) was used to compare the detected rate of alleles mismatch per sample and the probability of identification (PID: Probability of Identity) were calculated. The PID is the probability that two genotypes that are not identical happen to coincide, the sib-PID (sibling-PID) is the probability of assuming that two individuals have a blood relationship, and the HW-PID (Hardy-Weinberg-PID) is probability when assuming no blood relationship to each other. The actual PID is HW-PID > PID > sib-PID. Particularly in mammals, a sib-PID <0.01 is considered to be desirable (Waits et al. 2001). In this study, analysis by the software GENECAP version 1.4 was done by default setting. The “Match probability” was set to “0.05”, and “Method to use” setting was selected by analyzing “sib”.

## RESULTS AND DISCUSSION

### 1. Development of microsatellite (SSR) markers

As a result of pair-end sequencing by HiSeq X Ten, a total of 14,866,986,000 bp of contig (raw sequence data) was obtained, within which 14,726,967,300 bp (99.1%) was evaluated as being “clean data (correct rate: > 99%)”. Furthermore, as a result of the subsequent assembly, the sequences of 339,864,595 bp of the genomic DNA of *Chimarrogale platycephalus* were obtained. Among these, microsatellite sequences were detected at 11,727 sites. Among them, for 11,176 of the microsatellite sites, we were able to design appropriate primers (Fig. 3).

**Fig. 3.**
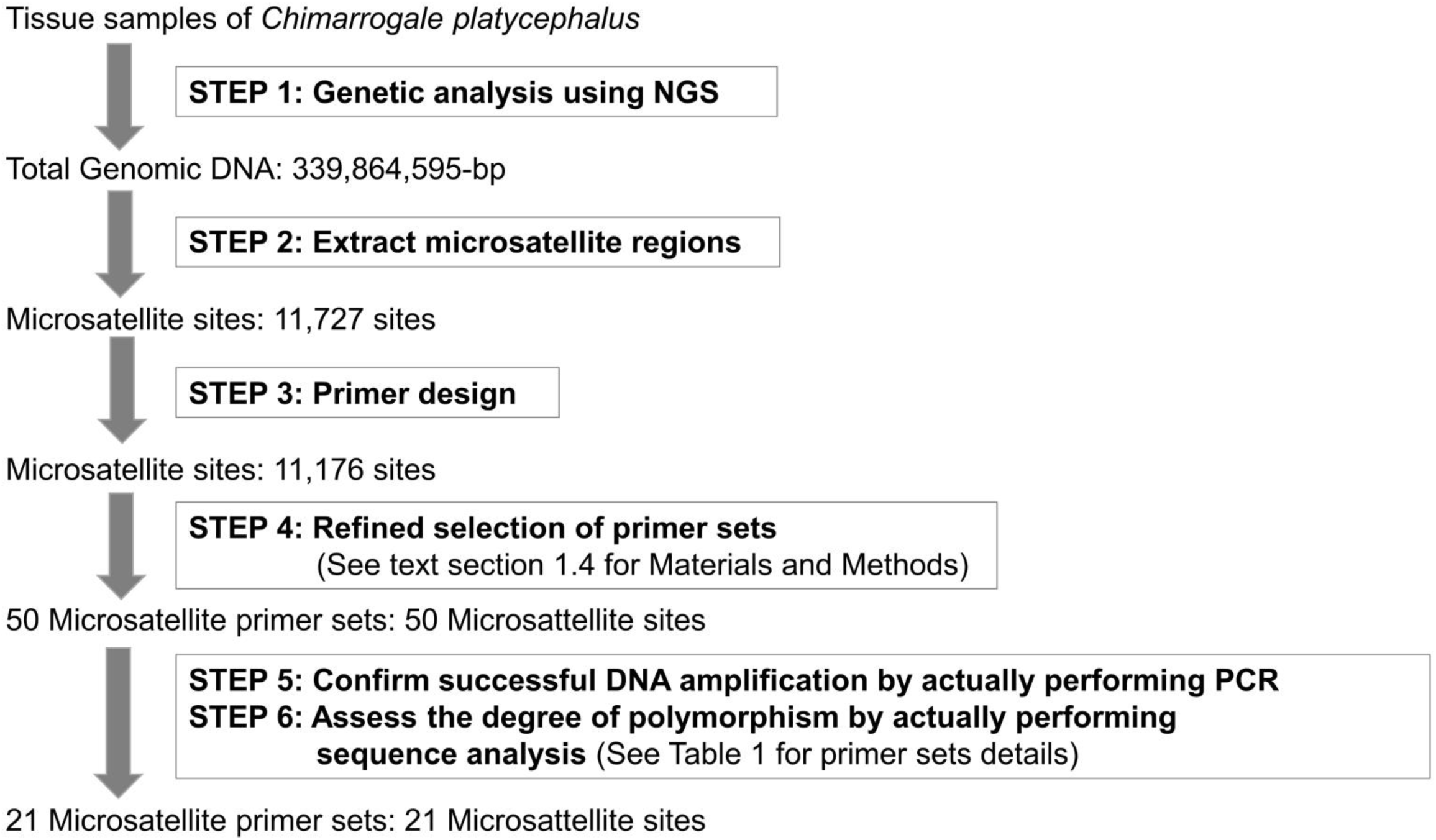
Research flow chart for developing microsatellite markers for a freshwater soricid shrew, *Chimarrogale platycephalus* (Soricomorpha, Soricidae), using a next generation sequencer (NGS). Please refer to the text for the details of the experiment at each step.

#### Selection of microsatellite sites for actual analyses

From the sequences containing the 11,176 microsatellites for which appropriate primers were designed, selection of target sites for our microsatellite analyses was performed based on the following four criteria: 1) Number of repeat units, 2) size of repeat units, 3) Tm value, and 4) specificity of the *C. platycephalus* sequence. As a result, 50 microsatellite markers were selected as candidates, PCR and subsequent electrophoresis were performed using these 50 primer sets, and it was confirmed whether DNA amplification could be performed without problems. With respect to the sequences in which DNA amplification could be confirmed in a series of experiments, DNA sequencing using the ABI 3130 xl auto-analyzer was conducted. As a result of this, 21 loci were considered to be useful as microsatellite markers. As a result of the sequencing of these 21 microsatellite sites of multiple *C. platycephalus* individuals, suitable genetic polymorphisms among the individuals were identified (Table 1).

**Table 1.**
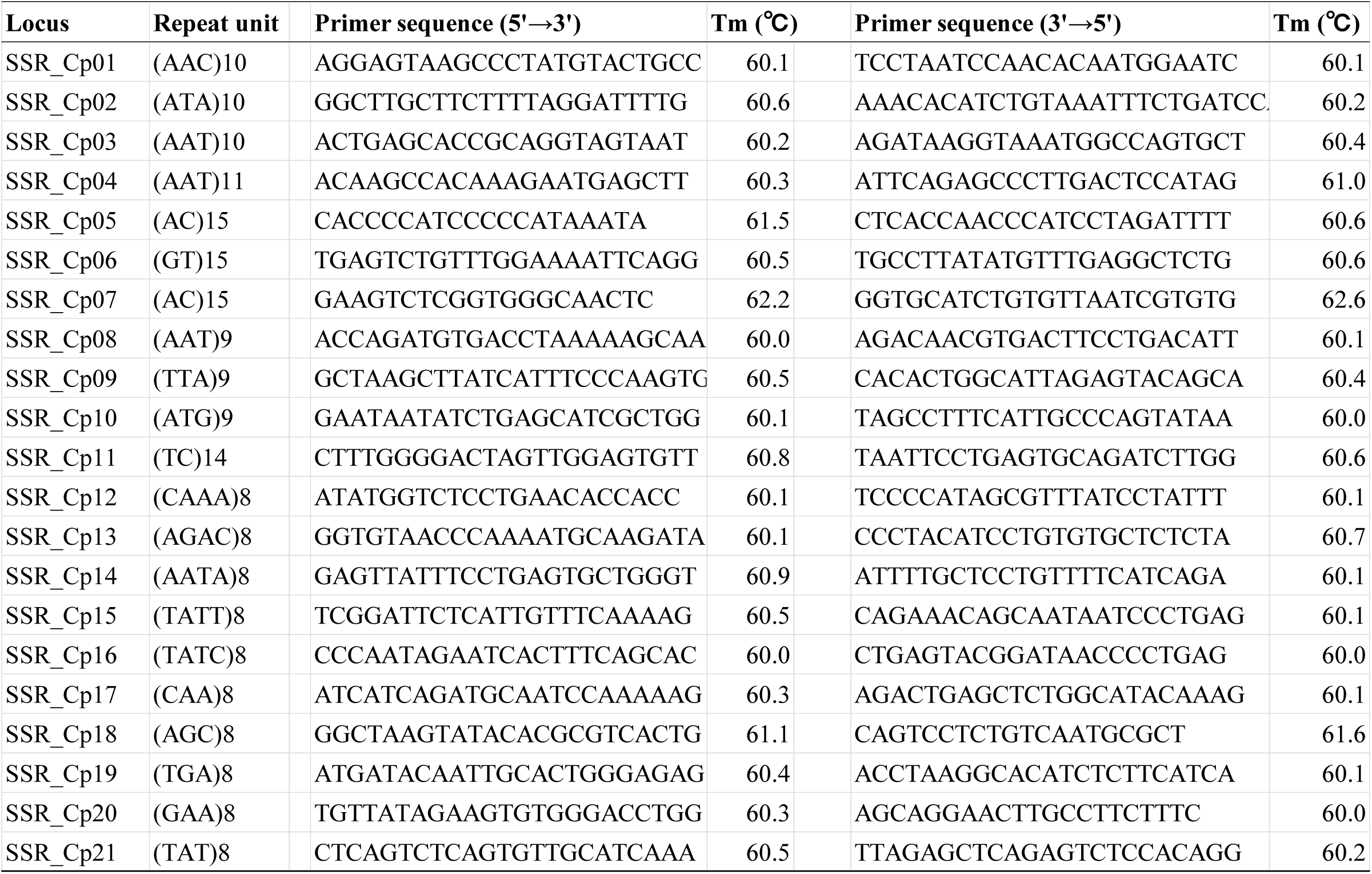
The 21 newly selected microsatellite markers used to examine the method of individual identification of the freshwater shrew, *Chimarrogale platycephalus* in this study, and their primer information.

### 2. Individual identification using the 21 microsatellite (SSR) markers

In this study, fecal specimens from individuals of *C. platycephalus* reared separately were used. As a result of collecting 29 fecal specimens from 12 *C. platycephalus* individuals and analyzing the 21 microsatellites’ sites for each of them, analysis was successful for 26 of the 29 specimens (Table 2). For the unsuccessful 3 samples, it was deemed that there were problems with the condition of the DNA sample of the feces. In general, DNA analysis from fecal specimens is difficult because the quantity DNA contained in the stool is very small. Also, analysis of nuclear DNA is particularly difficult, because the number of copies of nuclear DNA is smaller than that of mitochondrial DNA. Furthermore, in the feces of *C. platycephalus* excreted near the water area, hydrolysis of DNA is likely to occur, which is considered to make analysis more difficult. Under such circumstances, it can be considered that the successful analysis of 26 of the 29 specimens is of great significance.

**Table 2.** Analyzed microsatellite marker numbers for each fecal specimen of *Chimarrogale platycephalus*. In order to examine the pros and cons of analysis from fecal samples, when a reared individual would die, microsatellite analysis was performed using tissue specimens from the dead individual and compared with data from the fecal specimens. This table also contains the results of microsatellite analysis using these tissue specimens as data for comparative references.

| No. Individual (Symbol) | No. of each sample | Specimen type used for genetic analysis | No. of analyzed SSR loci |
| --- | --- | --- | --- |
| No. 01 (A) | A1 | F | 17 |
|  | A2 | F | * |
|  | A3 | T | 21 |
| No. 02 (K) | K1 | F | 19 |
|  | K2 | F | 19 |
|  | K3 | T | 21 |
| No. 03 (M) | M1 | F | 20 |
|  | M2 | F | 20 |
|  | M3 | F | * |
|  | M4 | T | 21 |
| No. 04 (S) | S1 | F | 16 |
|  | S2 | F | 16 |
|  | S3 | F | 20 |
| No. 05 (U) | U1 | F | 20 |
|  | U2 | F | 21 |
|  | U3 | F | * |
|  | U4 | T | 21 |
| No. 06 (Y) | Y1 | F | 20 |
|  | Y2 | F | 20 |
|  | Y3 | T | 21 |
| No. 07 (Z) | Z1 | F | 21 |
|  | Z2 | F | 21 |
|  | Z3 | F | 20 |
|  | Z4 | T | 21 |
| No. 08 (KR) | KR1 | F | 21 |
|  | KR2 | F | 21 |
| No. 09 (GM) | GM1 | F | 16 |
|  | GM2 | F | 21 |
| No. 10 (MT) | MT1 | F | 15 |
|  | MT2 | F | 18 |
|  | MT3 | F | 21 |
| No. 11 (MS) | MS1 | F | 18 |
|  | MS2 | F | 21 |
| No. 12 (WR) | WR1 | F | 20 |
|  | WR2 | F | 20 |
|  |  | T: Tissue | *Sample that could not be analyzed |
|  |  | F: Feces |  |

Even within those 26 fecal specimens that were able to be analyzed, there were some microsatellite sites that could not be analyzed. As such, although the number of loci analyzed from fecal samples varied from 15 to 21 loci, it was confirmed that for each of the sequenced gene loci analyzed, that sufficiently reliable results were obtained. As we will be described in detail later, our analyses were successful in providing sufficiently reliable data to allow individual identification even from fecal specimens of *C. platycephalus*. Six tissue samples were added to this, and analysis was performed with 32 samples in total.

#### Establishment of individual identification method using GENECAP

Microsatellite analysis from fecal specimens of different individuals identified different alleles depending on the locus (Table 3). However, no polymorphisms were found among the three markers (SSR_Cp10, Cp17 and Cp18). When all the analyzed loci were compared and examined, the probability of correct individual identification became sib-PID <0.01, using at least 12 loci. It is considered that the positive identification probability of mammals should be PID-sib <0.01 (Waits et al. 2001). In addition, the results of these microsatellite based analyses were 100% consistent with the actual excretion recorded by the examined individuals. Even in the case of using only 8 microsatellite sites, a sib-PID=0.0124 value was obtained, still providing relative degree of confidence in positive identification of individuals being possible. Probably, of the 21 microsatellite sites selected in this study, if 9-10 sites could be analyzed, individual identification may be considered possible (Table 4).

**Table 3.** Analysis results for each microsatellite marker.

| **Locus** | **No. of**<br>**analyzed samples** | **No. of**<br>**detected alleles** | **SSR size**<br>**range (bp)** |
| --- | --- | --- | --- |
| SSR_Cp1 | 31 | 2 | 167-173 |
| SSR_Cp2 | 28 | 5 | 169-193 |
| SSR_Cp3 | 28 | 5 | 177-195 |
| SSR_Cp4 | 32 | 3 | 165-171 |
| SSR_Cp5 | 32 | 7 | 103-122 |
| SSR_Cp6 | 25 | 9 | 151-183 |
| SSR_Cp7 | 32 | 6 | 123-161 |
| SSR_Cp8 | 32 | 9 | 117-141 |
| SSR_Cp9 | 31 | 5 | 148-164 |
| SSR_Cp10 | 29 | 1 | 166 |
| SSR_Cp11 | 23 | 9 | 166-198 |
| SSR_Cp12 | 32 | 4 | 150-166 |
| SSR_Cp13 | 31 | 5 | 98-114 |
| SSR_Cp14 | 30 | 7 | 168-192 |
| SSR_Cp15 | 26 | 8 | 158-198 |
| SSR_Cp16 | 31 | 7 | 155-179 |
| SSR_Cp17 | 32 | 1 | 142 |
| SSR_Cp18 | 32 | 1 | 143 |
| SSR_Cp19 | 29 | 2 | 178-181 |
| SSR_Cp20 | 31 | 2 | 136-139 |
| SSR_Cp21 | 31 | 8 | 170-194 |

**Table 4.** Microsatellite markers based individual identification validity results calculated using the software GENECAP version 1.4. The results are shown separately for cases in which the possibility of “allele drop-out” and “allele drop-in” could not be eliminated and were thus excluded subsequent analysis.

| Matches of samples |  | Sib-PID | HW-PID |
| --- | --- | --- | --- |
| **Case1** |  |  |  |
|  | A1, A3 | 1.28E-05 | 7.04E-16 |
|  | K1, K2, K3 | 5.97E-06 | 1.17E-18 |
|  | M2, M4 | 2.36E-06 | 4.18E-17 |
|  | U1, U2, U4 | 3.85E-06 | 2.54E-16 |
|  | Y1, Y3 | 8.85E-07 | 8.39E-19 |
|  | Z1, Z2, Z3, Z4 | 1.86E-06 | 6.49E-19 |
|  | MS1, MS2 | 2.49E-05 | 5.24E-14 |
|  | WR1, WR2 | 5.18E-07 | 1.41E-20 |
| **Case2** |  |  |  |
|  | M1, M2,M4 | 2.07E-06 | 2.78E-18 |
|  | S1, S2, S3 | 1.45E-03 | 3.53E-08 |
|  | Y1, Y2, Y3 | 4.62E-06 | 1.35E-16 |
|  | GM1, GM2 | 1.64E-04 | 1.85E-11 |
|  | KR1, KR2 | 1.10E-06 | 6.71E-18 |
|  | MT1, MT2, MT3 | 1.24E-02 | 1.94E-05 |
| Case 1: In case of non-exclude the allele drop-out/drop-in |  |  |  |
| Case 2: In case of exclude the allele drop-out/drop-in |  |  |  |

The *C. platycephalus* individuals, targeted for analysis in this study for individual identification, were all collected within relatively close regional populations, including some from the same mountain stream in Fukushima Prefecture, Japan (Fig. 4, Table5). Under even these challenging conditions, being able to identify the individual with an accuracy of 100% demonstrates the high accuracy results of these microsatellite markers. In addition, since it has been confirmed that these microsatellite markers are also effective between distant populations (e.g., Nagano Prefecture, Honshu, and Kumamoto Prefecture, Kyushu), these should be evaluated as highly versatile markers that are not limited to intraspecific lineages or area groups.

**Fig. 4.**
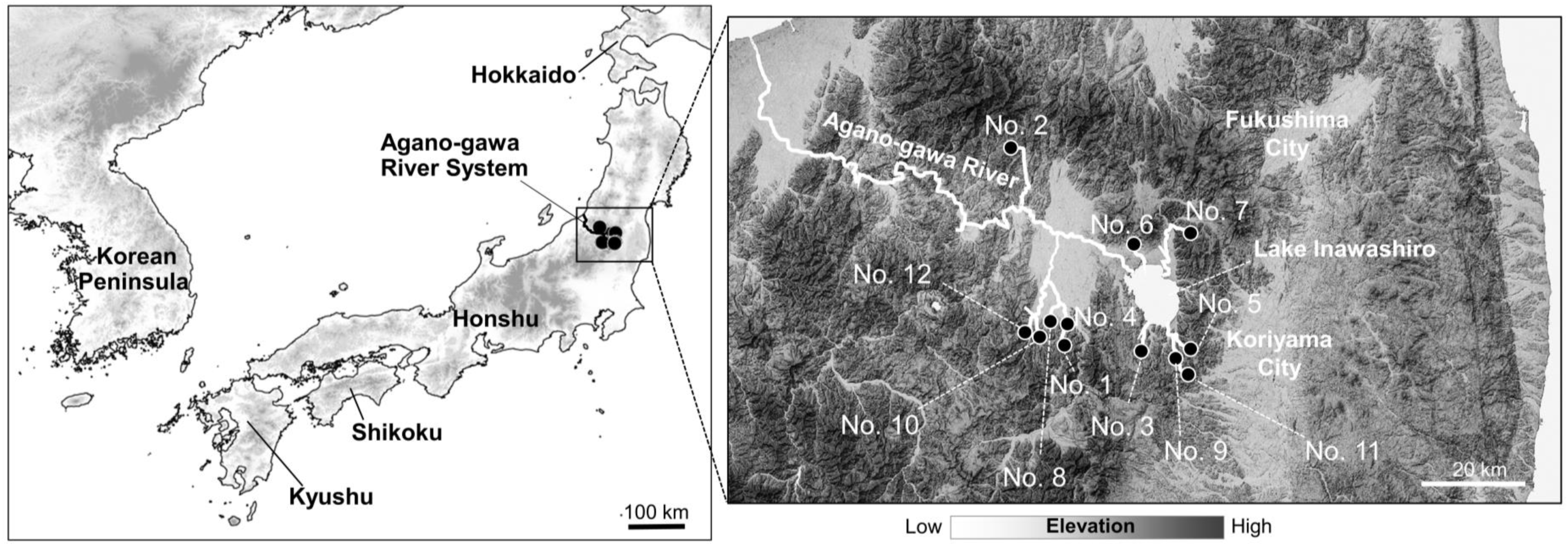
Collection sites of the twelve *Chimarrogale platycephalus* individuals used in the individual identification experiment, with the microsatellite markers developed in this study. In this study, 12 individuals collected from relatively close populations (Fukushima Prefecture) were analyzed for the purpose of verifying their individual identifiability. The numbers on the map (collected individual numbers) correspond to the individual numbers in Table 5.

In general, microsatellite analysis errors are considered to be likely to occur in cases where “allele dropout” has occurred (Kirov et al. 2000, Buchan et al. 2005). This is a phenomenon in which an apparent homozygous state occurs, because the amplification of one gene fails in spite of the fact that the allele is in a heterozygous state. In this study for the targeted fecal specimens of *C. platycephalus*, allele dropout detection was particularly pronounced. This tendency is considered to be caused by the extremely small amount of DNA contained in fecal specimens of *C. platycephalus*. However, in many specimens, by carrying out multiple repeated analyses, it was observed that an allele peak identified correctly in evaluation from homozygous to heterozygous in the same site. From this, it can be seen especially for microsatellite analysis of fecal specimens, that it is important to carry out repeated analyses several times in order to improve reliability by eliminating misidentified allele peaks. The samples which were able to analyze only small number of loci tended to have the “allele dropout” status, and in such cases, “allele dropout” with multiple loci was often observed. Regarding the samples, which were observed only homozygosity, “allele dropout” may have been detected. On the other hand, the “allele drop-in” phenomenon was observed only at one locus of one sample. In general, the “allele drop-in” is low in reproducibility and often has irregular waveforms in sequencing, so it can be judge relatively easily. In microsatellite analysis, at least in fecal samples, multiplex PCR should be avoided. Although there are some things to keep in mind when using fecal samples as such, effectiveness results will be obtained if we use with caution them.

**Table 5.** Details of each *Chimarrogale platycephalus* specimen’s information, used in their genetic analysis experiments: Identification numbers of reared individuals and symbol attached to reared individuals in the aquarium, collection dates and collection sites.

| **No. Individual (Symbol)** | **Collection Date** | **Collection Site** | **Prefecture** | **Latitude** | **Longitude** |
| --- | --- | --- | --- | --- | --- |
| No. 01 (A) | Nov. 15, 2017 | Onuma | Fukushima | 37.372 | 139.875 |
| No. 02 (K) | Nov. 10, 2017 | Kitakata | Fukushima | 37.757 | 139.766 |
| No. 03 (M) | Oct. 26, 2017 | Koriyama | Fukushima | 37.352 | 140.073 |
| No. 04 (S) | Nov. 15, 2017 | Onuma | Fukushima | 37.380 | 139.874 |
| No. 05 (U) | Sep. 12, 2017 | Koriyama | Fukushima | 37.369 | 140.171 |
| No. 06 (Y) | Nov. 10, 2016 | Yama | Fukushima | 37.574 | 140.040 |
| No. 07 (Z) | Nov. 5, 2016 | Inawashiro | Fukushima | 37.594 | 140.171 |
| No. 08 (GM) | Oct. 17, 2018 | Onuma | Fukushima | 37.377 | 139.839 |
| No. 09 (KR) | Oct. 22, 2018 | Koriyama | Fukushima | 37.356 | 140.148 |
| No. 10 (MT) | Oct. 17, 2018 | Onuma | Fukushima | 37.370 | 139.810 |
| No. 11 (MS) | Oct. 22, 2018 | Koriyama | Fukushima | 37.347 | 140.151 |
| No. 12 (WR) | Oct. 17, 2018 | Onuma | Fukushima | 37.375 | 139.806 |

The significance of this study is that identification of individuals from *C. platycephalus* feces is possible without using invasive means such as capture. It is considered that the microsatellite (SSR) markers developed in this study are able to contribute to the accumulation of fundamental ecological knowledge of *C. platycephalus*, and also the conservation of such endangered species.

## AUTHOR CONTRIBUTIONS

KT(K. Tojo), HY and TS conceived and designed the research. All the authors conducted with sampling. SN, KH and KT (K. Tokura) reared individually and collected feces. HY and TS conducted genetic analyses, and the subsequent development the microsatellite markers using the NGS. HY performed genotyping using SSR markers. All the authors discussed the composition of the paper, which was mainly written by KT (K. Tojo) and YH.

## ACKNOWLEDGEMENTS

Author contributions: We thank Mr. T. Motoki (EAC Cop., Matsumoto), Dr. A. Ichikawa (BOGA Ltd., Matsumoto), T. Suzuki, K. Yano (Shinshu University) for their cooperation with the field research and collection of specimens. This study was supported by the Japan Society for the Promotion of Science (JSPS) KAKENHI (JP23657064, and JP16K14807 to KT), grants from the River Environment Fund (2018-5211-015 to KT) of River and Watershed Environment Management, and a research grant from the Institute of Mountain Science, Shinshu University (KT).

